# Substrate-rigidity dependent migration of an idealized twitching bacterium

**DOI:** 10.1101/581348

**Authors:** Ahmet Nihat Simsek, Andrea Braeutigam, Matthias D. Koch, Joshua W. Shaevitz, Joshua W. Shaevitz, Yunfei Huang, Gerhard Gompper, Benedikt Sabass

## Abstract

Mechanical properties of the extracellular matrix are important determinants of cellular migration in diverse processes, such as immune response, wound healing, and cancer metastasis. Moreover, recent studies indicate that even bacterial surface colonization can depend on the mechanics of the substrate. Here, we focus on physical mechanisms that can give rise to substrate-rigidity dependent migration. We study a “twitcher”, a cell driven by extension-retraction cycles, to idealize bacteria and perhaps eukaryotic cells that employ a slip-stick mode of motion. The twitcher is asymmetric and always pulls itself forward at its front. Analytical calculations show that the migration speed of a twitcher depends non-linearly on substrate rigidity. For soft substrates, deformations do not lead to build-up of significant force and the migration speed is therefore determined by stochastic adhesion unbinding. For rigid substrates, forced adhesion rupture determines the migration speed. Depending on the force-sensitivity of front and rear adhesions, forced bond rupture implies an increase or a decrease of the migration speed. A requirement for the occurrence of rigidity-dependent stick-slip migration is a “sticky” substrate, with binding rates being an order of magnitude larger than unbinding rates in absence of force. Computer simulations show that small stall forces of the driving machinery lead to a reduced movement on high rigidities, regardless of force-sensitivities of bonds. The simulations also confirm the occurrence of rigidity-dependent migration speed in a generic model for slip-stick migration of cells.

## 1 Introduction

Surface migration is ubiquitous among both prokaryotic and eukaryotic cells. In the case of bacteria, various dedicated machineries for surface migration are known to facilitate nutrient search and surface colonization. Most notably, micrometer-sized filaments termed type-IV pili that are extended and retracted in a cyclic fashion, allow the bacteria to pull themselves forward. This mechanism of surface migration is common among mostly Gram-negative bacteria such as *Pseudomonas aeruginosa, Neisseria gonorrhoeaea, Escherichia coli*, and *Myxococcus xanthus*.^1–11^ Pilus-driven migration is termed “twitching” and visually resembles a random slip-stick motion.

Presently, it is an open question how the migration of bacteria depends on the substrate rigidity. Experimentally, substrate colonization by the pathogen *P. aeruginosa* and by *E. coli* appear to depend on the rigidity of various substrates,^12,13^ which may be relevant for host infection and for hygiene-related issues.^14–17^ A rigidity-dependent migration speed may result from physical effects related to the cell-substrate adhesion. In addition, bacterial migration is also regulated by surface adaptation processes that may involve mechanosensing. Different modes of bacterial mechanosensing have been suggested, in particular in the context of the transition from planktonic swimming to surface colonization. Putative mechanosensing pathways involve type-IV pili, pilus-related proteins, and outer membrane porins.^6,9,18–24^

In contrast to bacterial surface migration, migration of mammalian cells has been studied systematically to decipher its intricate dependence on environmental cues. Rigidity-dependent migration is for instance thought to be critical for the development of the nervous system,^25–27^ responses of the innate immune system,^28^ as well as cancer metastasis.^29–31^ An important hallmark of these physiological processes is the cellular adaptation to extracellular mechanics based on actomyosin cytoskeletal contractility, integrin engagement, and membrane dynamics. Thereby, eukaryotic cells can control their migration speed and direction depending on the mechanical rigidity of their substrate.^32,33^ Crawling cells forming pronounced substrate adhesions often undergo extension-contraction cycles, whereby elastic stress is built up and released. Such slip-stick-like motion is most clearly seen if the migration is constrained to a one-dimensional track.^34,35^

A prerequisite for the full understanding of complex biological cell-substrate interactions is the quantitative knowledge of the underlying physical mechanisms. Much work has been devoted to the theoretical understanding of how extracellular mechanics affects the migration of eukaryotic cells. Intricate models have been suggested, such as the cellular Potts model^36–38^, phase-field models^39–41^, and actin-flow and clutch models.^39,42–44^ Cells can also exhibit directed motion in gradients of rigidity, which is called durotaxis, and has recently sparked renewed theoretical interest.^45–48^ With regard to bacteria, mechanical modelling efforts have mostly focused on the understanding of pilus-based migration.^5,11,21,49,50^ However, the role of substrate rigidity for bacterial migration has not yet been studied.

Migration in phases of slip and stick is a distinctive physical characteristic found for bacteria and eukaryotic cells. Here, we explore the possibility that this slip-stick migration generically results in durokinesis, i.e., a dependence of the migration speed on substrate rigidity. To this end, we employ a highly idealized model twitcher. As a possible physical realization of a twitching cell, we have in mind the situation depicted in Fig. 1a,b. We assume a one-dimensional migration on a purely elastic gel with a tuneable rigidity. At the rear end, the cell body is anchored to the substrate via an elastic spring. At the front, an extending and retracting spring drives migration. This frontal driving mechanism represents, e.g., a bacterial pilus and naturally leads to a slip-stick motion. Binding and unbinding of the front and rear adhesions to the substrate, as well as the retraction of the front appendage are modeled stochastically. This model predicts stick-slip motion if the binding rates of the adhesions are an order of magnitude larger than the unbinding rates in absence of force. Analysis of this model further reveals a critical dependence of the migration speed on the force-sensitivity of the adhesion bonds. If the rear adhesions are more force sensitive, twitchers increase their migration speed as rigidity increases. Conversely, if the front adhesions are more force sensitive, the migration speed decreases with increasing substrate rigidity. Hence, durokinesis can be a result of physical mechanisms leading to a positive or a negative correlation of migration speed and rigidity of the extracellular environment.

**Fig. 1.**
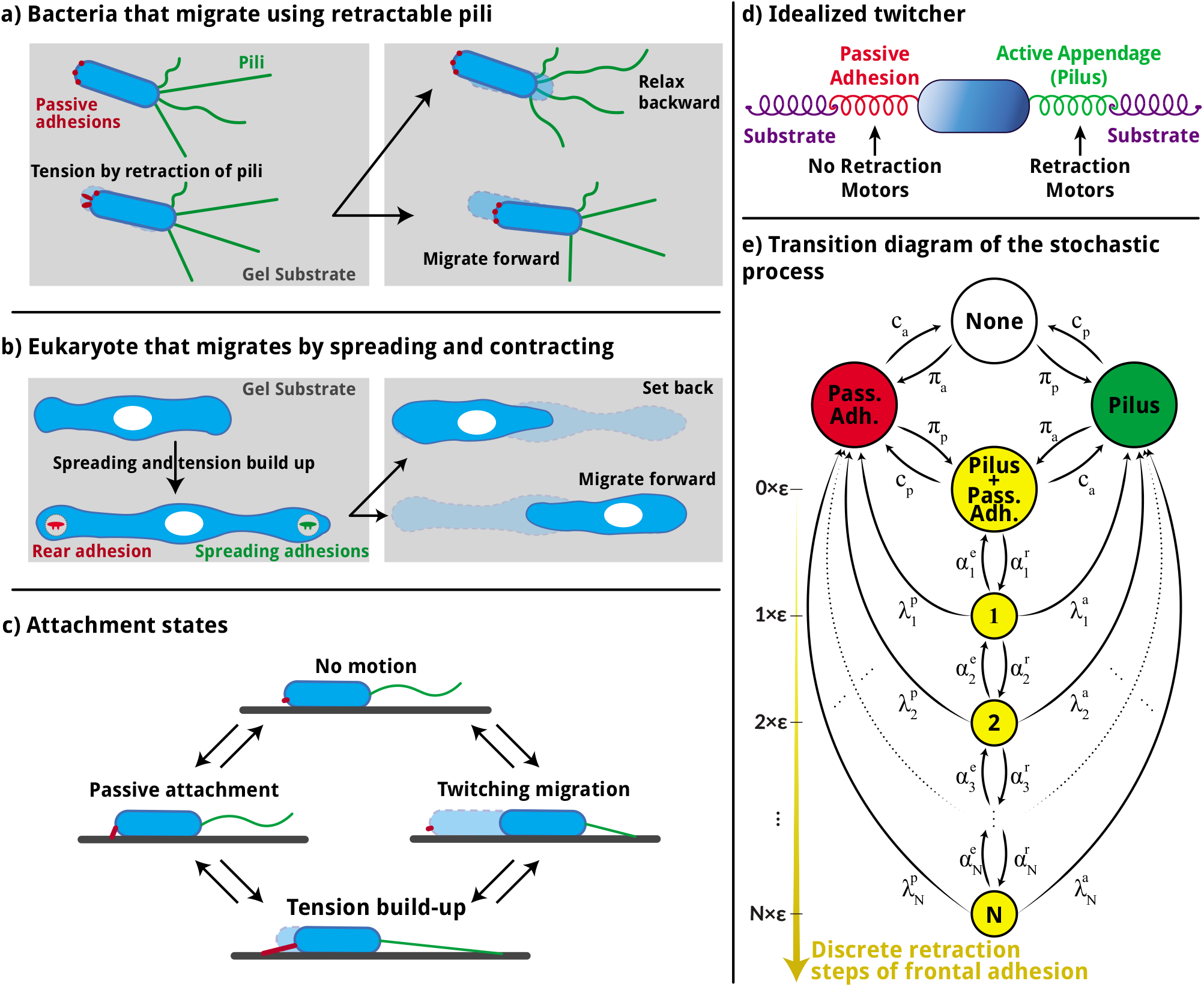
a) A bacterium moving by retractable pili. Passive adhesions are indicated at the rear of the bacteria. b) Eukaryotic cell moving through cyclic spread and contraction of its body. c) Stick-slip migration can be conceptually be divided into different states of substrate attachment. d) Sketch of the idealized model twitcher. e) State diagram of the analytical model for the twitcher. A “traffic light” color scheme is used to describe the motion in different states. None: the twitcher has no attachments to the substrate. Green: only the front appendage is attached and the twitcher migrates with constant speed. Red: only the passive adhesion is attached and no movement occurs. Yellow: both of the appendages are attached and the retraction builds up tension in discrete steps.

The article is structured as follows. In Sec. 2.1–2.3, we describe and solve a master equation for a twitcher model with discretized dynamics. In Sec. 2.4–2.7, we explore the biophysical predictions of the model and vary the system parameters. Sections 2.4 and 2.5 also contain a semi-quantitative discussion of the physical effects at work. The analytical work is complemented in Sec. 3 by simulations of a model with continuous mechanical stretch. These simulation results are compared to the discrete model and the effect of retraction motor stall force is investigated. Conclusions are presented in Sec. 4.

## 2 A master equation approach

We describe the dynamics of the twitcher with an analytically solvable master equation governing the substrate binding dynamics and mechanical stress in the twitcher.

### 2.1 Model and governing equations

The cell is modelled as a one-dimensional object. At the frontal, active side, an elastic spring can bind to the cell substrate and exert traction force. In the case of bacteria, this spring can be thought of as an idealized pilus. At the rear end, the cell attaches to the substrate via specific or unspecific passive adhesions that are again modeled as a spring connecting the cell body with the substrate, see Figs. 1c,d for schematic illustrations. In order to be able to construct an analytically solvable system, we assume that the tension in the system relaxes very quickly such that the balance of the elastic forces in the system occurs instantaneously. The cell migrates forward by retraction of its frontal appendage, which has an initial rest length of ℓ = ℓ_0_. Retraction and elongation of the frontal appendage are represented in a discretized form where the rest length ℓ of the frontal spring changes in small steps of length *ε*. With these assumptions, the system dynamics is described by transitions between the following states

- **S**_none_ - none of the appendages are attached to the substrate,
- **S**_adh_ - only the passive adhesion is attached to the substrate,
- **S**_pil_ - only the front appendage (pilus) is attached to the substrate,
- **S**_both_(or **S**_0_) - both appendages are attached to the substrate but there is no tension,
- **S**_1_ - both appendages are attached and the frontal appendage is retracted by *ε*,
- …
- **S**_N_ - both appendages are attached and the frontal appendage is retracted by *N ε*.

The state diagram of the stochastic process is given in Fig. 1e. In the state **S**_none_, the substrate is not bound and no motion can occur. Similarly, no motion occurs in the state **S**_adh_ where the cell is only anchored by its passive, rear adhesion. In the state **S**_pil_, the cell is only bound to the substrate via the active, frontal appendage. It is assumed that the cell moves here with constant average speed without build-up of elastic forces in the system. In state **S**_both_, both front and rear appendages are attached to the substrate and the front adhesion is fully extended with rest length ℓ_0_. From there on, retraction of the frontal appendage can occur through transitions to the states **S**_n_ with *n* ∈{0, 1, …, *N*}where the rest length of the front appendage is shortened as ℓ=ℓ_0_ − *nε*.

We next consider the force balance in one of the states **S**_n_ in which the front and rear adhesions are both bound to the substrate. The elastic stretch of the front appendage is denoted by 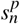 where the index *p* stands for “pilus”. The stretch of the passive adhesion is denoted by 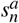 and the stretch of the substrate springs is denoted by 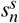. The sign of the stretches is chosen such that we can write the force balance among the frontal adhesion, the rear adhesion, and the substrate as

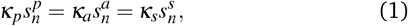

where *κ_p,a,s_* represent the spring constants of front adhesion (pilus), rear adhesion, and substrate, respectively. For simplicity, we assume that the springs modeling the substrate and the passive adhesion have a vanishing rest length. The rest length of the front adhesion is ℓ_0_ during initial attachment. Since the overall distance that is initially spanned by the serial springs does not change when the rest length changes from ℓ_0_ to ℓ, we have

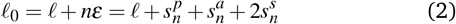

for all *n* ∈{0,1,…,*N*}. Combining Eqns. (1) and (2), the force acting across the system in state **S**_both_ is given by

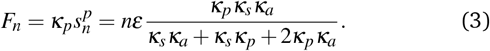

We next consider the transition rates between the states. Front and rear appendages are assumed to attach to the substrate with constant rates *π_p_* and *π_a_*, respectively. The corresponding detachment (rupture) rates depend on the applied force.^5,7,51,52^ Here, we mainly assume slip bonds where the rupture rate increases exponentially with the loading force. The slip-bond rupture rates for frontal and rear adhesion are given by

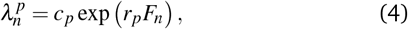

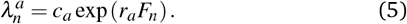

The retraction and elongation rates of the frontal appendage also assumed to be load-dependent and we choose

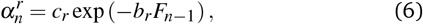

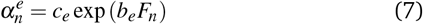

for retraction and elongation, respectively. This choice ensures that the speed of the retraction decreases as the force increases. Such a force-velocity relationship is a qualitative characteristic of both bacterial pili,^53^ and myosin-powered cellular contractions.^54^ After multiple retraction steps, the force increases to a point where the retraction stalls. The stall force is defined as the force at which average retraction speed vanishes and the retraction state at which stalling occurs is determined within our model by

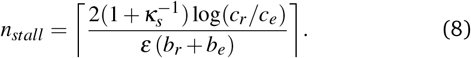

The stochastic processes can be summarized by the following master equations

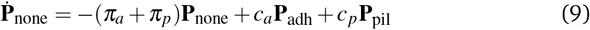

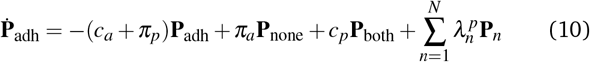

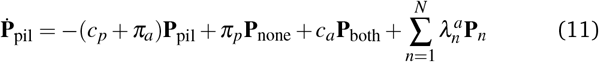

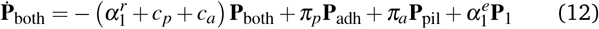

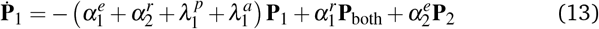

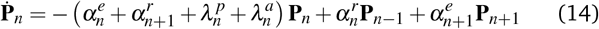

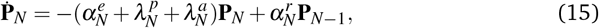

where **P**_*x*_ and 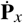 denotes the probability that the system is in the state **S**_*x*_ at time *t* and its time derivative, respectively. For simplicity, we will subsequently employ the notations 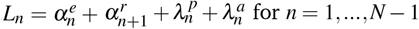 and 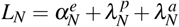.

### 2.2 Stationary solution of the master equation

We search for a stationary solution where 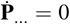 for all states. Observe from Eq. (15) that 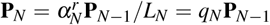, where *q_N_* is used to denote relevant transition rates in a compact form recursively. Substitution into Eq. (14) for *n*=*N*−1 and rearrangement gives

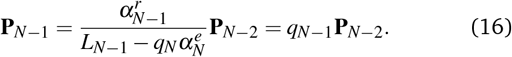

This process is continued until **P**_1_=*q*_1_**P**_both_ is obtained. Then, recursive substitution yields the relation

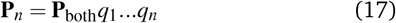

between any tension state **S**_*n*_ and **S**_both_. The resulting relations allow the elimination of **P**_*n*_ in Eqns. (10),(11), which then yield

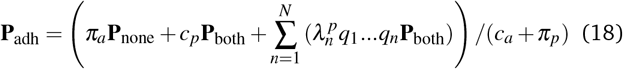

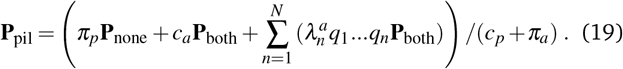

Substitution into equation (9) and reorganization gives

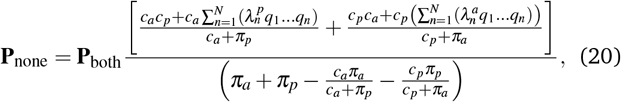

which also allows **P**_adh_ and **P**_pil_ to be expressed solely in terms of rates and **P**_both_. Finally, normalization of probabilities requires

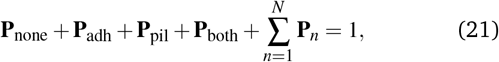

which is used to represent **P**_both_and all other probabilities only in terms of rates.

### 2.3 Calculation of the mean speed

The twitcher described above can move in two ways. First, through unhindered pilus retraction with speed *v_r_* in state **S**_pil_. Pilus retraction speed under zero load is given by *v_r_* =*ε*(*c_r_* – *c_e_*).

Second, the twitcher moves instantaneously after rupture of one bond from the tensed states **S**_*n*_ to relax the remaining attached elastic appendage. When the front adhesion ruptures, the bacterium returns to its original position. Thus, this event does not contribute to total migration. In contrast, rupture of the rear, passive adhesion in state **S**_*ñ*_ leads to a forward motion of the total retracted pilus length *ñε*. Therefore, the mean distance covered per unit time 〈*ẋ*〉, which we call the mean speed, is given by

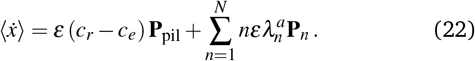

As shown in the appendix, the mean speed quickly converges for an increasing number of tensed states *N*, see Fig. 6. In the following, we choose *N*=5 if not mentioned otherwise. The employed parameters are listed in Table 1.

**Table 1.**
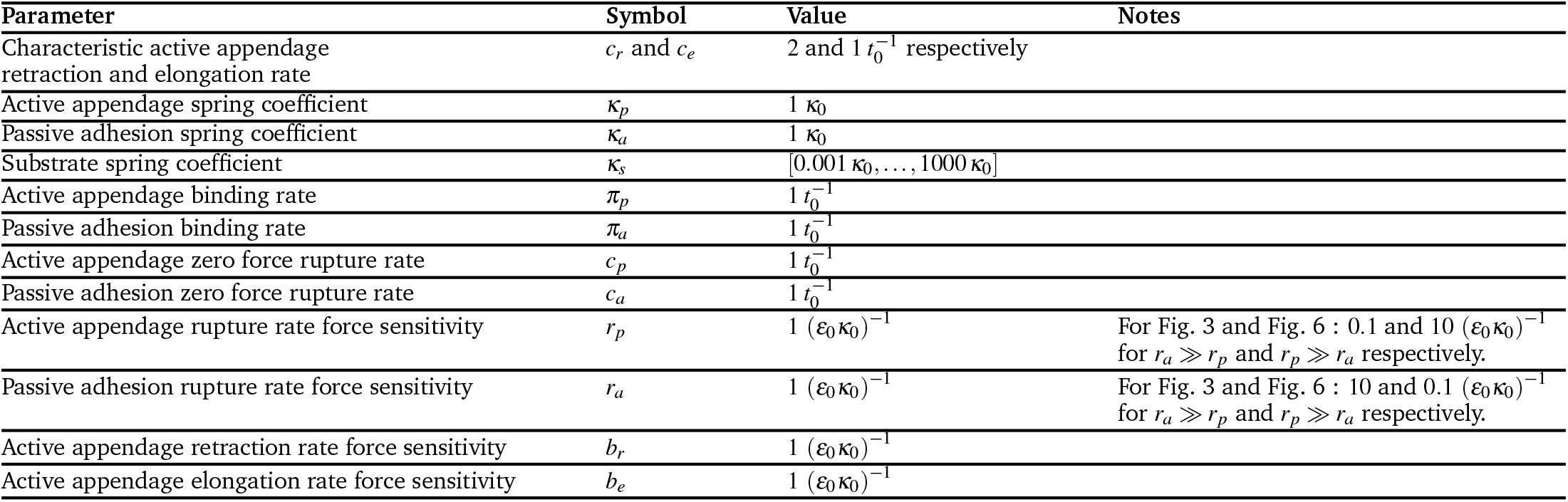
Parameters for the analytical discrete-state model. The time scale is *t*_0_, the length scale is *ε*_0_, and the stiffness scale is *κ*_0_.

### 2.4 Rigidity-dependent migration: at a first glance

We investigate the dependence of the mean speed 〈*ẋ*〉 on the substrate rigidity *κ_s_*. The solution of the master equation leads to lengthy analytical expressions for the speed, too long to be presented here. The expression for the case *N* = 1 is given in the Appendix. We consider qualitative features of the results. Figure 2 shows the effect of substrate rigidity on the mean speed for different force sensitivity parameters, *r_a_* and *r_p_*, of the rupturing bonds. The curves show a non-linear relation between substrate rigidity and migration distance. For *r_a_* > *r_p_*, the rear adhesion ruptures faster than the front adhesion under force and the mean speed increases with substrate rigidity. Conversely, *r_a_* < *r_p_* implies that the front adhesion ruptures faster and the mean speed decreases on increasing rigidity.

**Fig. 2.**
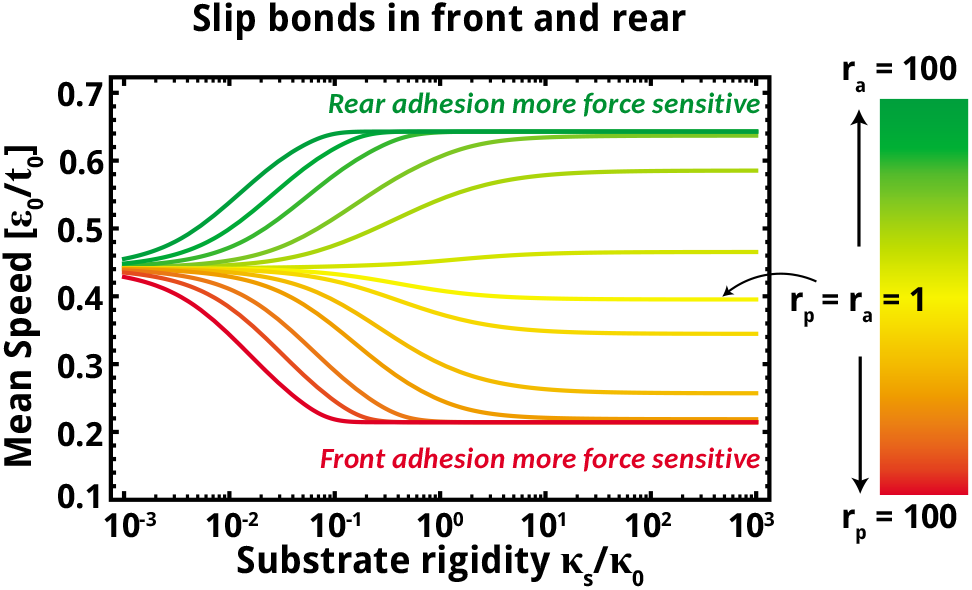
Effect of substrate rigidity on mean twitching speed. Yellow lines correspond to the case that front and rear bonds have equal force-sensitivity *r_a_* = *r_p_* = 1. Red color tones correspond to *r_a_* < *r_p_* with *r_a_* = 1, green color tones correspond to *r_a_* > *r_p_* with with *r_p_* = 1. Note that the migration speed increases or decreases with substrate rigidity, depending on on the force-dependent rupture rates. The varied force sensitivity parameters are taken from the set {1,2,5,10,25,50,100}. Front and rear adhesions are assumed to be slip bonds. See Table 1 for the other parameters and the non-dimensionalization.

### 2.5 Rigidity-dependent migration: analysis and limiting cases

In order to gain a qualitative interpretation of the rigidity-dependence of speed, we consider Eq. (3) determining the force on the bonds. A simplification occurs if the spring constants of the front and rear adhesions are set to the same value *κ_p_* = *κ_a_* ≡ *κ_i_*, which we refer to as internal spring constant characterizing the internal rigidity. Thus, we have

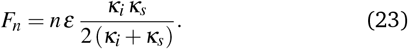

Figure 3a illustrates the role of the internal spring constant *κ_i_* for substrate-dependent twitching migration. The mean speed of a very stiff twitcher with *κ_i_* ≫ 1/(*εr_a,b_*) changes strongly with substrate rigidity. For small internal rigidity, the effect of substrate rigidity on twitching speed vanishes almost completely.

**Fig. 3.**
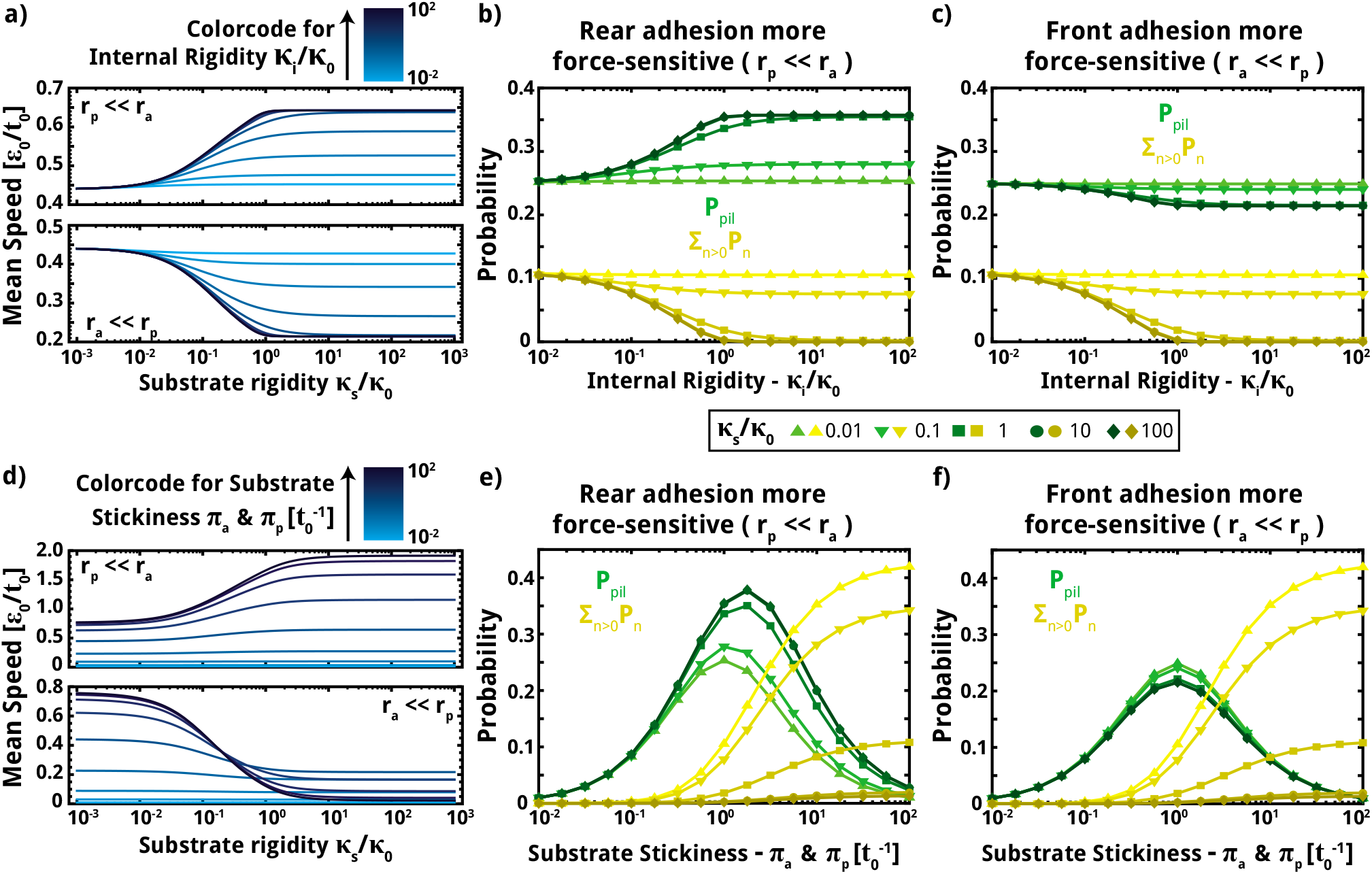
a-c) Role of the internal twitcher rigidity *κ_i_* for durokinesis. The impact of external substrate rigidity on the mean speed increases with internal rigidity. This result is due to a change in state probabilities illustrated in b-c). Increasing *κ_i_* shifts the balance between states in which the twitcher can be tensed, 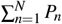, and the state where it retracts the frontal adhesion freely, *P*_pil_. d-f) The substrate binding rates *π_p_* and *π_a_* have a strong impact on the migration dynamics. The state probabilities in e-f) show that for small binding rates, *π_p,a_* < *c_p,a_*, the state probability *P*_pil_ dominates, which implies that migration occurs predominantly through unhindered, but rare, retraction of the frontal appendage. For *π_p,a_* ~ *c_p,a_ P*_pil_ still dominates but both motion are observed more often. On “sticky” substrates with *π_p,a_* ≫ *c_p,a_*, the states probabilities 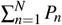 dominate, which implies rigidity-dependent stick-slip motion. Parameters in a) and d) are logarithmically spaced with *κ_i_/κ*_0_ and 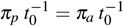 being in the range 10^−2^,…,10^2^. See Table 1 for the other parameters.

We now assume that *κ_i_* has a fixed, finite value and consider limiting cases for the substrate rigidity. On substrates with vanishing rigidity, we have lim_*κ_s_*→0_*F_n_* =0. Retraction of the appendages on soft substrates does not produce much force. Consequently, the detachment rates of frontal and rear appendages are given by 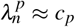 and 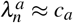, leading to a migration speed that is independent of rupture mechanics. Accordingly, twitchers with different force-dependent rupture rates have the same mean speed on very soft substrates, as seen in Fig. 2. On substrates with higher rigidity, the forces *F_n_* exceed the typical scales 1/*r_a,b_* and therefore start to affect the twitching dynamics through the bond rupture rates, Eqns. (4) and (5). For very rigid substrates, we have lim_*κ*_*s*→∞__*F_n_* = *nεκ_i_*/2. The elastic contraction within the cell dominates here. Hence substrate rigidity plays no role for the rupture rates and mean speeds become independent of *κ_s_*, as seen in Fig. 2.

To obtain a simplified expression that qualitatively captures the rigidity-dependence of the mean speed, we consider a model with only one retraction state, *N* = 1 and assume that the binding and unbinding rate constants are equal, *c_p_* = *c_a_* = *π_a_* = *π_p_* = 1. For this case, the analytical solution simplifies to

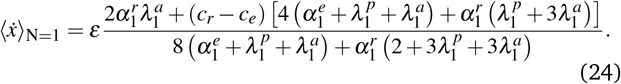

The numerator contains two terms, where the first results from relaxation events after a contracted state and the second results from motion in the state **S**_pil_ where only the front appendage is bound. To see how these terms are affected by substrate rigidity, we consider the case in which the bond at front is weaker, *r_a_* ≪ *r_p_*, and assume for simplicity that the rigidity of the twitcher is very high *κ_i_* ≫ *κ_s_*. Thus, the rupture rates are given by 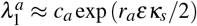 and 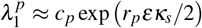. Large substrate rigidities allow the retraction to build up significant forces, i.e., *r_p_εκ_s_*/2 ≫ 1, and eventually lead to 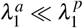. This relation implies that the first term in the numerator of Eq. (24) vanishes. Hence, twitchers with *r_a_* ≪ *r_p_* migrate slower on rigid substrates. In the opposite case that the bond at the passive, rear adhesion is weaker, *r_a_* ≫ *r_p_*, both terms in the numerator of Eq. (24) remain important, allowing for an increase in speed on more rigid substrates.

### 2.6 Mean speed and state occupation probabilities

The mean speed depends on rigidity via its effect on the state occupation probabilities that directly appear in the mean-speed equation (22). Figures 3b-c show these occupation probabilities **P**_pil_ and 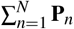 for the two cases of twitchers slowing down on higher rigidity *r_p_* ≫ *r_a_* and twitchers speeding up on higher rigidity *r_p_* ≫ *r_a_*. An increase in internal rigidity decreases the probability to be in the tensed states 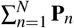. Instead, the system spends more time in states with unilateral attachment at the front or at the rear. For *r_p_* ≪ *r_a_*, high internal rigidity leads to a high probability for unilateral adhesion only at the front, **P**_pil_. In this state, unhindered retraction of the front appendage produces fast migration. For *r_p_* ≫ *r_a_*, high internal rigidity implies that the system spends more time in the state with only the rear appendage attached, consequently, migration speed is small.

Figure 3d illustrates the impact of “substrate stickiness”, i.e., the binding rates *π_p_* and *π_a_*, on the migration dynamics. As seen, low binding rates *π_p,a_* ≪ *c_p,a_*, do not permit adhesion of the twitcher and therefore the mean speed vanishes. In the regime of intermediate stickiness *π_p,a_* ~ *c_p,a_*, the motion is dominated by the state **S**_pil_ where the twitcher experiences no resistance, see Fig. 3e-f. Therefore, tensed states are suppressed and the substrate rigidity plays a reduced role. In contrast, high binding rates, *π_p,a_* ≫ *c_p,a_*, imply that both appendages are frequently attached to the substrate, leading to frequent build-up of contractile tension, i.e., the mean speed is dominated by slip-stick events resulting from bond rupture, see Fig. 3e-f. Thus, rigidity-dependent migration based on slip-stick motion requires a sticky substrate.

### 2.7 Effect of catch-bond dynamics

The above-employed slip-bond model is an important modeling paradigm.^51,55–58^ However, the lifefetime of certain bonds is known to increase for intermediate mechanical loads, which is called a catch bond.^7,52,59^ A prominent example are bacterial type I pili encoded in the *fimH* gene.^60^To studymotion of twitchers with catch bond adhesions, we employ the rupture rates

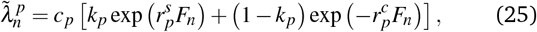

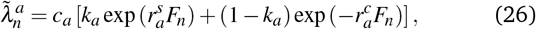

for the front and rear adhesion respectively. In Fig. 4a-top the pilus has a catch-bond constant *k_p_* = 0.025 and the passive adhesion is a slip bond. The data shows that a pilus with catch-bond mechanism still causes an increase of the mean speed with substrate rigidity, however, a maximum of mean speed can appear for intermediate substrate rigidity. This maximum increases and shifts towards softer rigidities as the catch force sensitivity 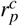 increases. Note that here the mean speed difference between the softest and hardest substrates is smaller than with the corresponding data for slip bonds. In Fig. 4a-bottom, the rear adhesion is a catch bond with *k_a_* = 0.025 while the pilus forms a slip bond with the substrate. Here, mean speed decreases as expected from the data for slip bonds, see Fig 2. However, compared to slip bonds, a catch bond can result in a sharper decrease of mean speed with increase of substrate rigidity. Finally, Fig. 4c shows exemplary results for twitchers having catch bonds at the front and rear. Here, increasing substrate rigidity reduces the mean speed and a global minimum can occur for intermediate rigidity.

**Fig. 4.**
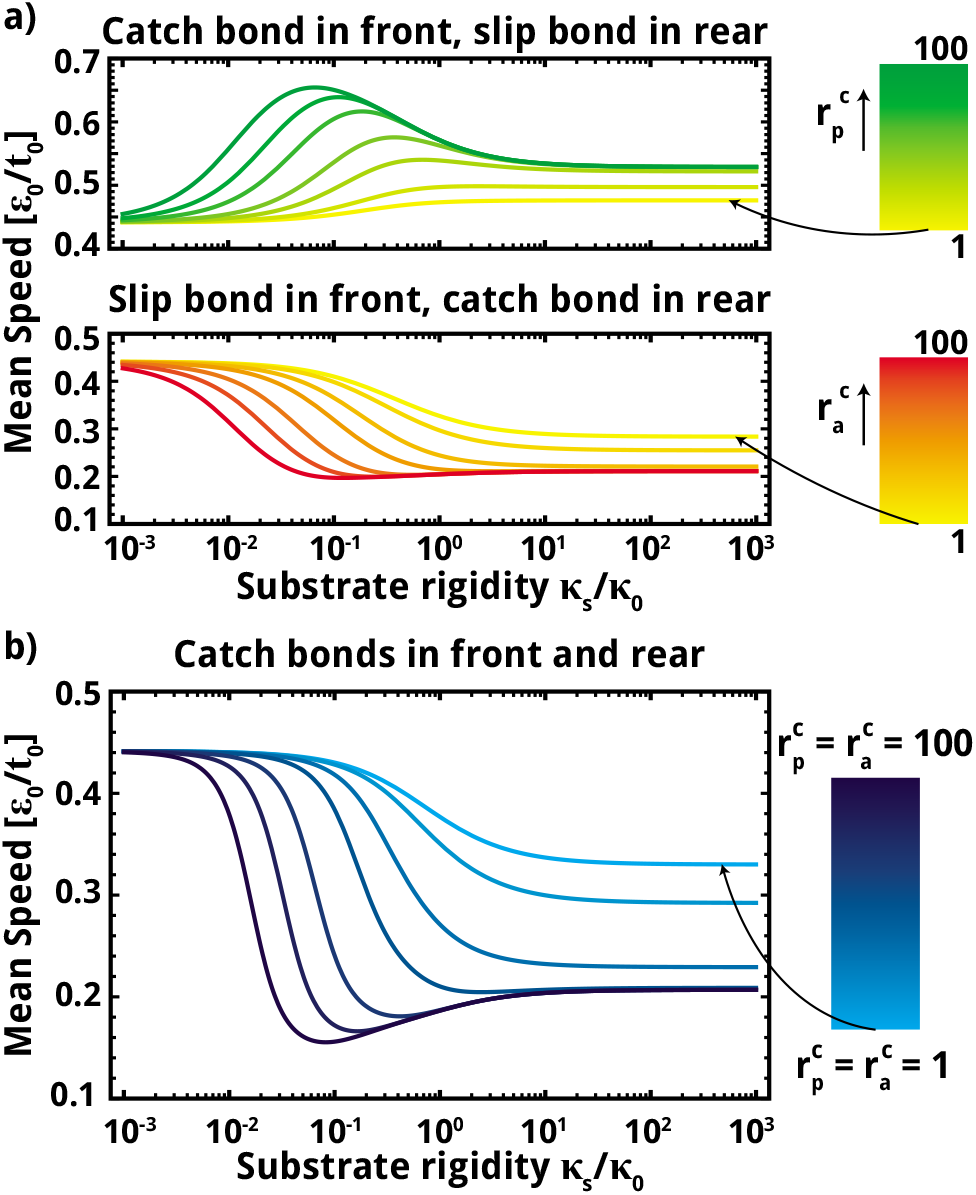
Effect of catch-bond dynamics on durokinesis of the twitcher. a) Top: Front adhesion is assumed to be a catch bond and the rear adhesion is a slip bond, 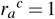. Bottom: Front adhesion is a slip bond and rear adhesion a catch bond, 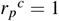. b) Both of the adhesions are assumed to be catch bonds. For all figures, the varied force sensitivity parameters are taken from the set {1,2,5,10,25,50,100}. See Table 1 for the other parameters.

## 3 Simulation of a twitcher model with continuous space variable

As an alternative approach to the discrete model studied in the previous section, we study a model in which the space variables and the elastic stretch are represented as continuous variables. This model corroborates the results from the discrete-state model and yields further insights regarding the distributions of migration speed and the role of the stall force.

### 3.1 Model and simulation method

We again consider the idealized twitcher shown in Fig. 1d. Now, however, the stretch of the front appendage *s_p_*, the rear appendage *s_a_*, and substrate *s_s_* are continuous variables. The state diagram and respective transition rates are shown in Fig. 5a. On introducing a viscous friction coefficient *η* and the instantaneous twitcher velocity *ẋ*(*t*), the continuous force balance reads

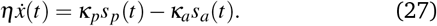

**Fig. 5.**
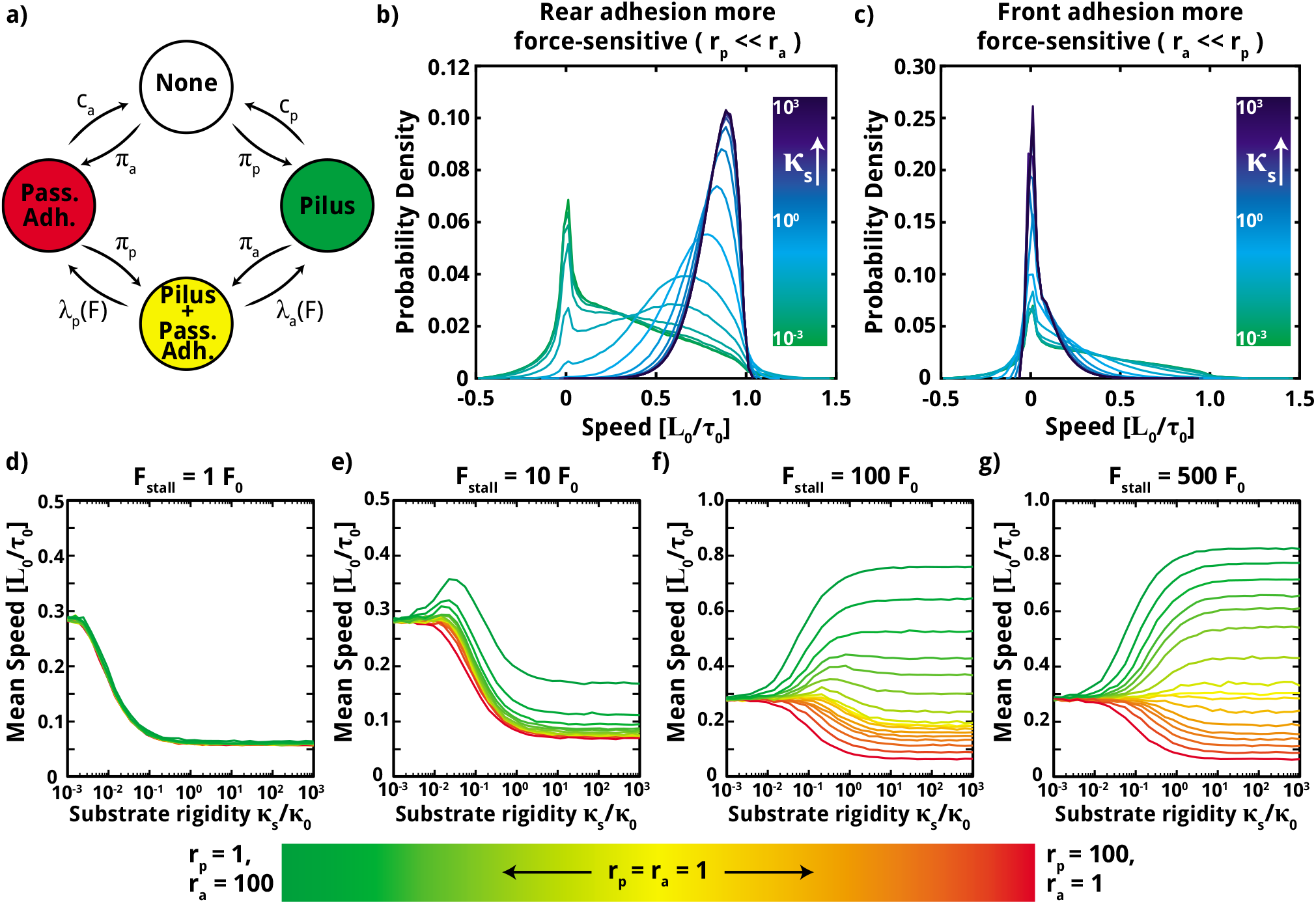
Stochastic simulation of a twitcher with continuous space variables. a) State diagram of the stochastic twitching process. Continuous retraction occurs in the states where the frontal adhesion (pilus) is bound. b-c) Distributions of coarse-grained migration speed. If the rear adhesion is rarely attached, migration speed is close to the retraction speed of the frontal appendage *v*_0_=1 *L*_0_/τ_0_. The statistics were obtained from from 100 simulation runs for each condition. d-f) Effect of retraction stall force on durokinesis of the twitcher. Low stall forces result in a decrease of mean speed on higher rigidity. Graphs in d) illustrate that this speed decrease is largely independent of the otherwise important parameters *r_a_* and *r_p_* since all plots fall on the same line. In f-g) large stall forces *F_stall_* ≥ 100 *F*_0_ imply a rare occurrence of stalling events. Here, the rigidity-dependence of the mean speed agrees qualitatively with the results from the discrete-state model.

Retraction of the front spring (the pilus) is governed by a linear force-velocity relationship that approximates experimental results as

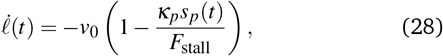

where *v*_0_ is the characteristic retraction speed in the absence of load.^5,51,61^ This force-velocity relationship implies that the retraction stalls when the load reaches *F*_stall_. We also assume that once the front spring is fully retracted, it detaches and is immediately replaced by a fully extended, unbound spring. Front and rear adhesions are assumed to be slip bonds with the detachment rates *λ_p_*(*s_p_*) = *c_p_*exp(*r_p_s_p_κ_p_*) and *λ_a_*(*s_a_*) = *c_a_*exp(*r_a_s_a_κ_a_*), as used above.

For the simulations, we employ a time scale τ_0_ and time discretization of step size Δ*t* = 10^−3^ τ_0_. All simulations are performed for a duration of 10000 τ_0_. In each time step, we draw a random number *R*_1_ from the uniform distribution of real numbers in the unit interval. The condition for the occurrence of a binding or rupture event in this timestep is then 1 − exp (−Δ*t*(*k*_1_ + *k*_2_)) > *R*_1_ where *k*_1,2_ are the rates of the possible events in the current state. If this condition is satisfied, a second random number is drawn from the uniform distribution and the condition *R*_2_ < *k*_1_/(*k*_1_ + *k*_2_) is checked for event selection. If the condition is satisfied, the event with rate *k*_1_ is chosen, else, the other event is chosen. Finally, the stretch states of the springs are propagated with Eqns. (27), (28). For comparison with the discrete-state model discussed in Sec. 2, we focus on the limit of fast relaxation with *η* = 0. Results for finite *η* are in qualitative agreement with the fast relaxation case, see Fig. 7 in the appendix.

### 3.2 Simulation results

Figure 5b-c shows speed distributions where the speed is calculated with a moving average of 10^3^ simulation steps, corresponding to τ_0_. For *r_a_* ≫ *r_p_*, an increase in substrate rigidity allows bacteria to move more freely as the passive adhesion ruptures more often. Hence, the speed distribution shifts towards the maximum retraction speed *v*_0_ = *L*_0_/τ_0_. In the case *r_p_* ≫ *r_a_*, the speed distributions shift to lower values as rigidity increases, since the passive adhesion can withstand larger forces than the active frontal adhesion. Also, the missing peak at speeds around *v*_0_ shows that the system is rarely in the state **S**_pil_ where only the front adhesion is bound. These findings agree qualitatively with the results of the discrete-state model in Sec. 2 which, however, only yields mean speeds.

Next, we study the influence of the stall force *F*_Stall_ on the migration speed. Figure 5d displays the mean speed of twitchers with very low stall force, meaning that the retraction of the front appendage stops a very small load. Since the stall prevents the build-up of significant elastic forces in the system, the adhesion bonds can survive for a long time. Nevertheless, some migration occurs on soft substrates where the weak forces can contract the substrate and move after rupture of the bonds. However, the mean speed decreases on more rigid substrates, regardless of order of the bond force-sensitivity parameters *r_a_* and *r_p_*. Figures 5e-g show that an increase of the stall force allows for an increase in mean speed on high rigidities. The stall force for the simulation data shown in Fig. 5g corresponds to the stall force employed in the analytical discrete-state model studied in Sec. 2. Here, the simulations results qualitatively agree very well with the results from the discrete-state model, compare Fig. 5g with Fig. 2.

## 4 Summary and concluding discussion

We propose a mathematical model to investigate the condition under which the physics of cellular slip-stick motion naturally leads to rigidity-dependent migration speed. The model is primarily inspired by bacterial twitching motion, but may also be of some value for describing slip-stick processes in mammalian cells.^43,62^ The twitcher is described as a one-dimensional body with one passive appendage at its rear and one appendage that can be extended and retracted at its front. Migration of the twitcher can occur in two modes: first, forward motion can occur through retraction of the front appendage if the rear appendage is not attached, second, jumps in the forward or backward direction can occur when bond rupture releases the elastic stress that the twitcher builds up in states where front and rear are both attached. Due to the simplifying description of the twitcher, migration can be studied with analytical approaches or simulated with standard algorithms.

In the analytical approach, we discretize the retraction and mechanical stretch of the twitcher to obtain a system of master equations. This model reveals a critical dependence of the migration dynamics on the force-sensitivity of the adhesion bonds. For substrates that are much more rigid than the internal rigidity scale, twitchers migrate faster if their rear adhesion bond is more sensitive to force. However, this effect vanishes on soft substrates, where large deformation of the substrate prevents the build-up of large forces during migration. Importantly, the occurrence of stick-slip motion is predicated on the binding rates being an order of magnitude larger than the rupture rates without tension. Hence, rigidity-dependent slip-stick requires a “sticky” substrate.

In order to confirm the results from the discrete-state model, we perform stochastic simulations of a continuum model. In the simulations, the contraction of the twitcher is modeled as a continuous process and a viscous relaxation timescale is introduces to avoid the assumption of instantaneous relaxation after bond rupture. Results from the simulations confirm the conclusions drawn from the discrete-state model. Moreover, the simulations reveal how stalling of the retraction of the front appendage can hinder movement. For stall forces that are much less than the typical forces produced during a typical retraction phase, the twitcher does not contract appreciably and the migration speed always decreases for increasing substrate rigidity. For large stall forces, the mean migration speed is similar to the one calculated in the discrete-state model and the force-sensitivity of the bonds determines whether the mean speed increases or decreases on higher rigidity.

Based on the results from the idealized model studied here, we expect that the suggested mechanism for durokinesis is generic for various cells that move in a slip-stick fashion. However, the biological complexities associated with migration of real bacteria or mammalian cells may modify or mask the mechanism studied here. With regard to bacterial migration, complicating effects are for instance the spatial arrangement of pili and other adhesions, complex adhesion bond dynamics, and active mechanosensing or adaptation. Such effects can be integrated in simulations of specific organisms. Finally, quantitative experiments are required for a comprehensive understanding of slip-stick migration of bacteria and mammalian cells on substrates of different rigidity.

## Conflicts of interest

There are no conflicts to declare.

## Acknowledgements

ANS acknowledges support by the International Helmholtz Research School of Biophysics and Soft Matter (IHRS BioSoft).

## Appendix

### Full expression of the mean speed for the case *N* = 1

Assuming only one retraction state, *N* =1, the mean speed can be given in full and reads

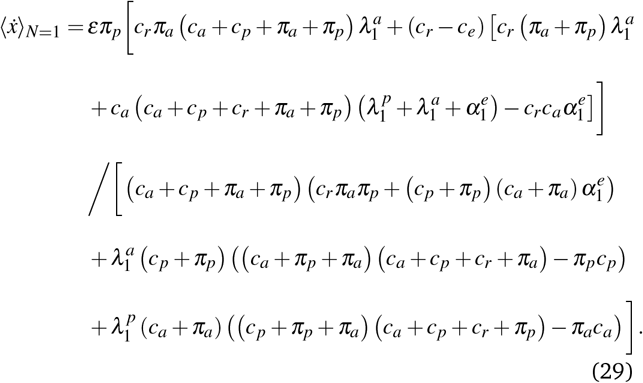

This equation corresponds to Eq. (24) given in Sec. 2.5 for the special case of *c_p_* = *c_a_* = *π_a_* = *π_p_* = 1.

### Role of the retraction state number *N* for the analytical calculations

Ideally, *N* would be chosen to be very large, thus allowing the retraction steps *ε* to represent motion on a molecular scale. However, the complexity of the algebraic solution for the mean speed increases drastically with the retraction state number *N*. Moreover, we find that the mean speed quickly converges for increasing *N* in our range of system parameters. For the parameter values in Table 1, we calculated the mean speeds on a range of substrate rigidities *κ_s_* as we vary maximum tension state *N* and compared the results in the Fig. 6. Based on theses results, *N* =5 is chosen as an approximation to limit the computational effort.

**Fig. 6.**
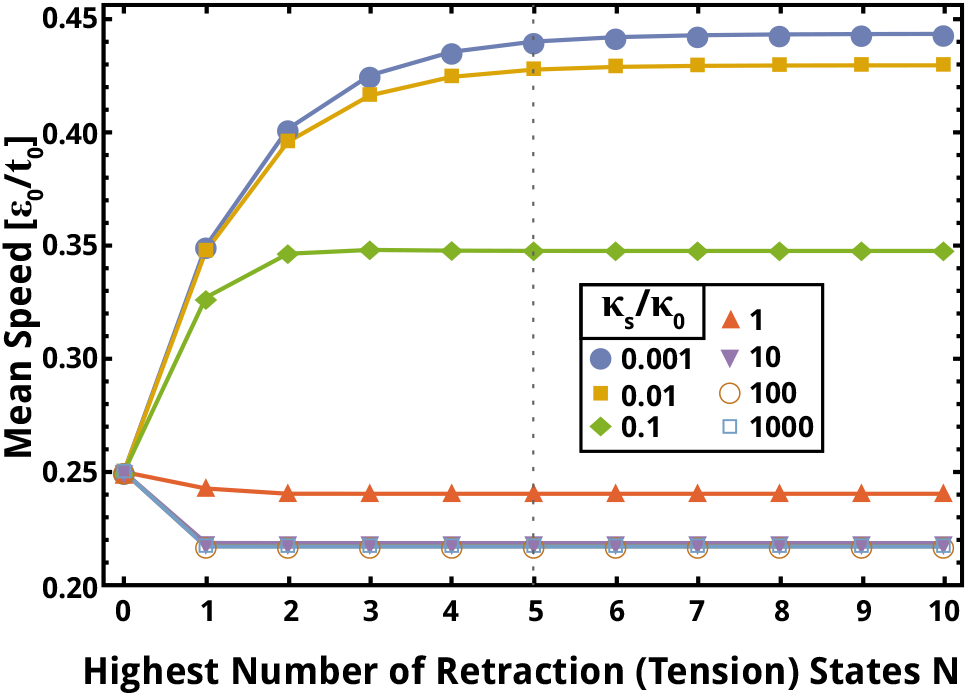
Comparison of mean speeds calculated with the discrete-state model for different number of maximum tension (retraction) states *N*. The mean speed quickly converges for an increasing number of tensed states *N*. For the data shown in the main text we chose *N* = 5. See Table 1 for other parameter values.

### Effect of a finite viscous relaxation time

Simulation results obtained with a finite friction coefficient *η* are shown in Fig. 7 together with the corresponding results assuming instantaneous fast relaxation. The dashed lines represent mean speeds for finite *η*, full lines represent the fast relaxation case. The plots demonstrate that the two cases are in qualitative agreement with each other. Finite relaxation times only play a role for *η* ≫ *κ*_0_τ_0_ and mainly affect the migration speed on very soft substrates. Hence, we expect that our results for fast relaxation also hold qualitatively for finite *η*.

**Fig. 7.**
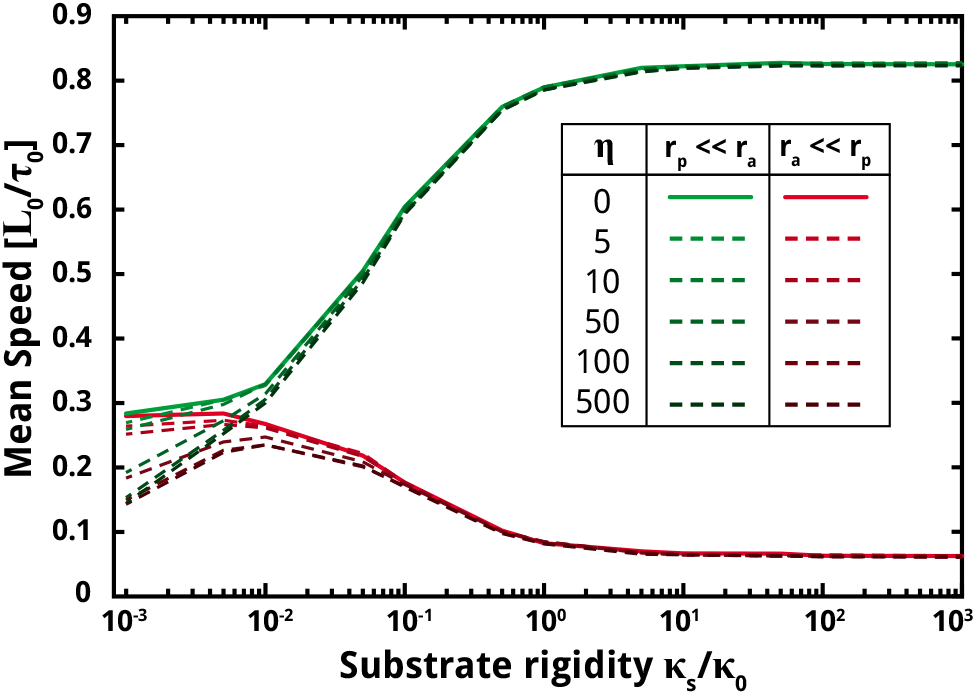
Simulations results for fast elastic relaxation with *η* = 0 and for finite viscous relaxation times with *η* > 0. Darker shades represent larger values of *η*. Simulations were performed using a time step Δ*t* = 10^−5^ τ_0_ where unit of *η* is 1(Δ*t*/τ_0_) *κ*_0_τ_0_. See Table 2 for the other parameter values.

### Parameter tables

The parameters employed for the analytical discrete-state model and the stochastic simulations are listed in Table 1 and Table 2 respectively. Corresponding numerical values are given in non-dimensional units. Unless stated otherwise, these values are used for the calculations and simulations.

**Table 2.**
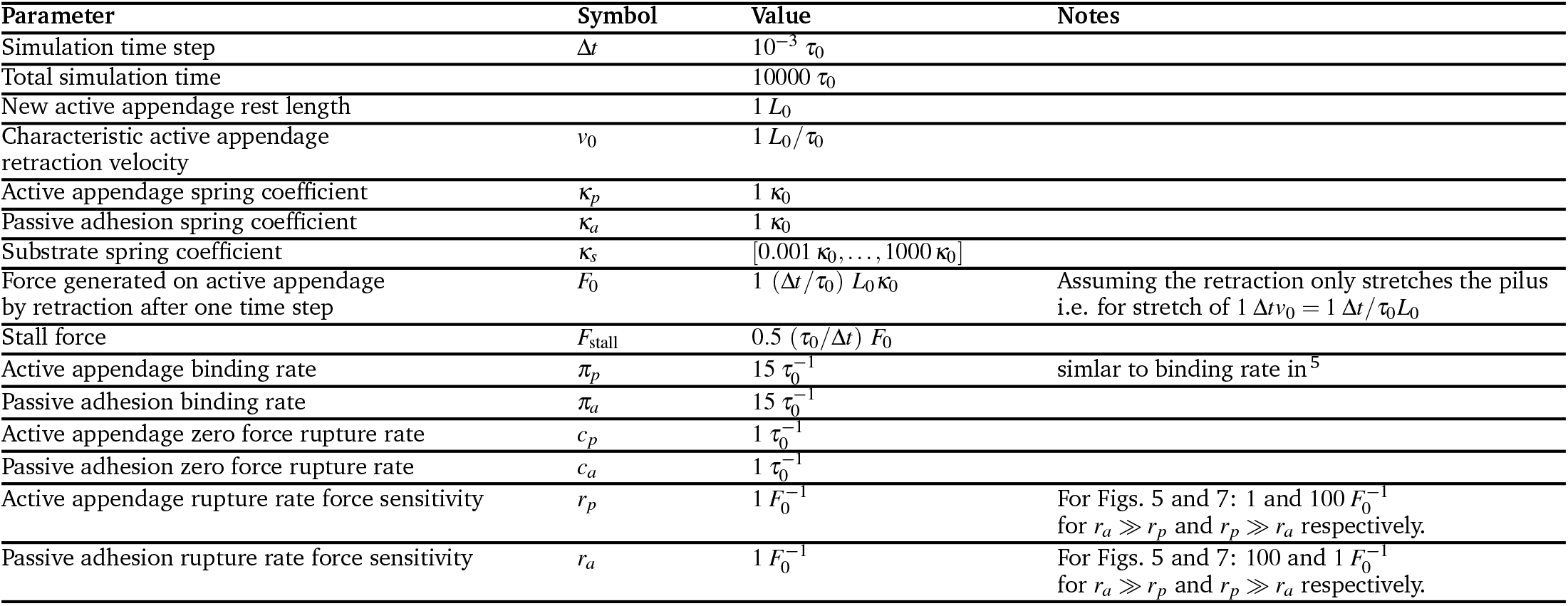
Parameters for the stochastic simulation. The time scale is τ_0_ = 1 sec, the length scale is *L*_0_, and the stiffness scale is *κ*_0_.

In Table 3, we provide numerical values for various physical quantities that were considered while developing the models. This table is meant to provide relevant units and scales for comparison to biological systems.

**Table 3.** Estimate of the parameters relevant for twitching Gram-negative bacteria.

| Parameter | Value [Range] | Unit | Notes |
| --- | --- | --- | --- |
| Relevant time scale | $[0.1, 120]$ | sec | see<sup>3-5</sup> |
| Relevant length scale | $[0.01, 10]$ | $\mu\text{m}$ | |
| Example type IV pili retraction/elongation speed | $[0.1, 10]$ | $\mu\text{m}/\text{sec}$ | see<sup>3-5</sup> |
| Example type IV pili stall force | $[80, 200]$ | pN | see<sup>5,57,61</sup> |
| Example type IV pili unbinding forces | $[1, 200]$ | pN | see<sup>5,57,61</sup> |
| Example type IV pili unbinding time scales | $[0.01, 1]$ | sec | see<sup>4,5</sup> |
| Example type IV pili binding rate | $(0, 20]$ | $\text{sec}^{-1}$ | see<sup>5</sup> |
| Example type IV pili stiffness | $2000 \pm 500$ | $\text{pN}/\mu\text{m}$ | see<sup>57</sup> |

## References

1 J. Henrichsen, Bacteriological reviews, 1973, 36, 478–503.

2 B. Maier and G. C. L. Wong, Trends in Microbiology, 2015, 23, 775–788.

3 B. Maier, Soft Matter, 2013, 9, 5667–5671.

4 J. M. Skerker and H. C. Berg, Proceedings of the National Academy of Sciences, 2001, 98, 6901–6904.

5 R. Marathe, C. Meel, N. Schmidt, L. Dewenter, R. Kurre, L. Greune, M. A. Schmidt, M. Mueller, R. Lipowsky, B. Maier and S. Klumpp, Nature communications, 2014, 5, 3759.

6 A. Persat, F. Yuki, J. N Engel, H. A Stone and Z. Gitai, Proceedings of the National Academy of Sciences of the United States of America, 2015, 112, 7563–7568.

7 A. Persat, C. Nadell, M. Kevin Kim, F. Ingremeau, A. Siryaporn, K. Drescher, N. Wingreen, B. Bassler, Z. Gitai and H. A Stone, Cell, 2015, 161, 988–997.

8 B. Sabass, M. D. Koch, G. Liu, H. A. Stone and J. W. Shaevitz, Proceedings of the National Academy of Sciences, 2017, 114, 7266–7271.

9 A. Yang, W. Shing Tang, T. Si and J. Tang, Biophysical Journal, 2017, 112, 1462–1471.

10 E. Miller, T. Garcia, S. Hultgren and A. Oberhauser, Biophysical journal, 2006, 91, 3848–56.

11 V. Zaburdaev, N. Biais, M. Schmiedeberg, J. Eriksson, A.-B. Jonsson, M. P Sheetz and D. A Weitz, Biophysical journal, 2014, 107, 1523–1531.

12 K. Kolewe, J. Zhu, N. R. Mako, S. Nonnenmann and J. Schiffman, ACS Applied Materials and Interfaces, 2018, 10, 2275–2281.

13 J. A Lichter, M. Todd Thompson, M. Delgadillo, T. Nishikawa, M.F Rubner and K. Van Vliet, Biomacromolecules, 2008, 9, 1571–8.

14 A. J Booth, R. Hadley, A. Cornett, A. A Dreffs, S. A Matthes, J. L Tsui, K. Weiss, J. Horowitz, V. Fiore, T. Barker, B. B Moore, F. Martinez, L. E Niklason and E. S White, American journal of respiratory and critical care medicine, 2012, 186, 866–876.

15 A. Verma, D. Bhani, V. Tomar, R. Bachhiwal and S. Yadav, Journal Of Clinical And Diagnostic Research, 2016, 10, 1–3.

16 F. Song and D. Ren, Langmuir: the ACS journal of surfaces and colloids, 2014, 30, 10354 – 10362.

17 N. Saha, C. Monge, V. Dulong, C. Picart and K. Glinel, Biomacromolecules, 2013, 14, 520 – 528.

18 K. Cowles and Z. Gitai, Molecular microbiology, 2010, 76, 1411–26.

19 Y. Luo, K. Zhao, A. E. Baker, S. L. Kuchma, K. A. Coggan, M. C. Wolfgang, G. C. Wong and G. A. OâĂŹToole, MBio, 2015, 6, e02456–14.

20 C. K. Ellison, J. Kan, R. S. Dillard, D. T. Kysela, A. Ducret, C. Berne, C. M. Hampton, Z. Ke, E. R. Wright, N. Biais, A. B. Dalia and Y. V. Brun, Science, 2017, 358, 535–538.

21 Y. Brill-Karniely, F. Jin, G. Wong, D. Frenkel and J. Dobnikar, Scientific Reports, 2017, 7, 45467.

22 A. Siryaporn, S. L Kuchma, G. A O’Toole and Z. Gitai, Proceedings of the National Academy of Sciences of the United States of America, 2014, 111, 16860–16865.

23 C. A. Rodesney, B. Roman, N. Dhamani, B. J. Cooley, P. Katira, A. Touhami and V. D. Gordon, Proceedings of the National Academy of Sciences, 2017, 114, 5906–5911.

24 C. K. Lee, J. de Anda, A. E. Baker, R. R. Bennett, Y. Luo, E. Y. Lee, J. A. Keefe, J. S. Helali, J. Ma, K. Zhao et al., Proceedings of the National Academy of Sciences, 2018, 115, 4471–4476.

25 L. A. Flanagan, Y.-E. Ju, B. Marg, M. Osterfield and P. A. Janmey, Neuroreport, 2002, 13, 2411.

26 L. Bollmann, D. Koser, R. Shahapure, H. Gautier, G. Holzapfel, G. Scarcelli, M. Gather, E. Ulbricht and K. Franze, Frontiers in Cellular Neuroscience, 2015, 9, 363.

27 K. Franze, P. A. Janmey and J. Guck, Annual review of biomedical engineering, 2013, 15, 227–251.

28 J. T. Mandeville, M. A. Lawson and F. R. Maxfield, Journal of leukocyte biology, 1997, 61, 188–200.

29 M. J. Paszek, N. Zahir, K. R. Johnson, J. N. Lakins, G. I. Rozenberg, A. Gefen, C. A. Reinhart-King, S. S. Margulies, M. Dembo, D. Boettiger et al., Cancer cell, 2005, 8, 241–254.

30 A. G. Clark and D. M. Vignjevic, Current opinion in cell biology, 2015, 36, 13–22.

31 D. Lachowski, E. Cortes, D. Pink, A. Chronopoulos, S. Karim, J. P Morton and A. del Rio Hernandez, Scientific Reports, 2017, 7, 2506.

32 J. Solon, I. Levental, K. Sengupta, P. Georges and P. Janmey, Biophysical journal, 2008, 93, 4453–61.

33 C. M Lo, H. B Wang, M. Dembo and W. Yl, Biophysical journal, 2000, 79, 144–52.

34 P. Maiuri, E. Terriac, P. Paul-Gilloteaux, T. Vignaud, K. McNally, J. Onuffer, K. Thorn, P. A. Nguyen, N. Georgoulia, D. Soong et al., Current Biology, 2012, 22, R673–R675.

35 K. Hennig, I. Wang, P. Moreau, L. Valon, S. De Beco, M. Coppey, Y. Miroshnikova, C. A. Rizo, C. Favard, R. Voituriez et al., bioRxiv, 2018, 354696.

36 P. J. Albert and U. S. Schwarz, Biophysical journal, 2014, 106, 2340–2352.

37 R. Allena, M. Scianna and L. Preziosi, Mathematical biosciences, 2016, 275, 57–70.

38 A. Goychuk, D. B. Brückner, A. W. Holle, J. P. Spatz, C. P. Broedersz and E. Frey, arXiv preprint arXiv:1808.00314, 2018.

39 E. Kuusela and W. Alt, Journal of mathematical biology, 2009, 58, 135.

40 D. Shao, H. Levine and W.-J. Rappel, Proceedings of the National Academy of Sciences, 2012, 109, 6851–6856.

41 F. Ziebert and I. S. Aranson, npj Computational Materials, 2016, 2, 16019.

42 P. DiMilla, K. Barbee and D. Lauffenburger, Biophysical journal, 1991, 60, 15–37.

43 K. A. Lazopoulos and D. Stamenović, Journal of biomechanics, 2008, 41, 1289–1294.

44 B. L. Bangasser, G. A. Shamsan, C. E. Chan, K. N. Opoku, E. Tüzel, B. W. Schlichtmann, J. A. Kasim, B. J. Fuller, B. R. McCullough, S. S. Rosenfeld et al., Nature communications, 2017, 8, 15313.

45 E. A. Novikova, M. Raab, D. E. Discher and C. Storm, Physical review letters, 2017, 118, 078103.

46 G. Yu, J. Feng, H. Man and H. Levine, Physical Review E, 2017, 96, 010402.

47 A.-R. Hassan, T. Biel and T. Kim, bioRxiv, 2018, 460170.

48 C. R. Doering, X. Mao and L. M. Sander, arXiv preprint arXiv:1806.00502, 2018.

49 W. T. Kranz, A. Gelimson, K. Zhao, G. C. Wong and R. Golestanian, Physical review letters, 2016, 117, 038101.

50 R. Groβmann, F. Peruani and M. Bär, New Journal of Physics, 2016, 18, 043009.

51 W. Pönisch, C. Weber, G. Juckeland, N. Biais and V. Zaburdaev, New Journal of Physics, 2017, 19, 015003.

52 S. Lecuyer, R. Rusconi, Y. Shen, A. Forsyth, H. Vlamakis, R. Kolter and H. A Stone, Biophysical journal, 2011, 100, 341–50.

53 B. Maier, M. Koomey and M. P. Sheetz, Proceedings of the National Academy of Sciences, 2004, 101, 10961–10966.

54 C. G. Galbraith, K. M. Yamada and M. P. Sheetz, J Cell Biol, 2002, 159, 695–705.

55 G. I. Bell, Science, 1978, 200, 618–627.

56 B. Sabass and U. S. Schwarz, Journal of Physics: Condensed Matter, 2010, 22, 194112.

57 A. Beaussart, A. E. Baker, S. L. Kuchma, S. El-Kirat-Chatel, G. A. O’Toole and Y. F. Dufrêne, ACS Nano, 2014, 8, 10723–10733.

58 A. Touhami, M. H Jericho, J. Boyd and T. J Beveridge, Journal of bacteriology, 2006, 188, 370–377.

59 W. Thomas, V. Vogel and E. Sokurenko, Annual review of biophysics, 2008, 37, 399–416.

60 M. M. Sauer, R. Jakob, J. Eras, S. Baday, D. Eriş, G. Navarra, S. Berneche, B. Ernst, T. Maier and R. Glockshuber, Nature Communications, 2016, 7, 10738.

61 B. Maier, L. Potter, M. So, C. D Long, H. S Seifert and M. P Sheetz, Proceedings of the National Academy of Sciences of the United States of America, 2003, 99, 16012–7.

62 C. E. Chan and D. J. Odde, Science, 2008, 322, 1687–1691.

